# Metagenomic shotgun analyses reveal complex patterns of intra- and interspecific variation in the intestinal microbiomes of codfishes

**DOI:** 10.1101/864439

**Authors:** Even Sannes Riiser, Thomas H.A. Haverkamp, Srinidhi Varadharajan, Ørnulf Borgan, Kjetill S. Jakobsen, Sissel Jentoft, Bastiaan Star

## Abstract

The relative importance of host-specific selection or environmental factors in determining the composition of the intestinal microbiome in wild vertebrates remains poorly understood. Here, we use metagenomic shotgun sequencing of individual specimens to compare the intra- and interspecific variation of intestinal microbiome communities in two ecotypes (NEAC and NCC) of Atlantic cod (*Gadus morhua*) – that have distinct behavior and habitats– and three *Gadidae* species that occupy a range of ecological niches. Interestingly, we find significantly diverged microbiomes amongst the two Atlantic cod ecotypes. Interspecific patterns of variation are more variable, with significantly diverged communities for most species’ comparisons, apart from the comparison between coastal cod (NCC) and Norway pout (*Trisopterus esmarkii*), whose community compositions are not significantly diverged. The absence of consistent species-specific microbiomes suggests that external environmental factors, such as temperature, diet or a combination there-off comprise major drivers of the intestinal community composition of codfishes.

**Importance:** The composition of the intestinal microbial community associated with teleost fish is influenced by a diversity of factors, ranging from internal factors (such as host-specific selection) to external factors (such as niche occupation). These factors are often difficult to separate, as differences in niche occupation (e.g. diet, temperature or salinity) may correlate with distinct evolutionary trajectories. Here, we investigate four gadoid species with contrasting levels of evolutionary separation and niche occupation. Using metagenomic shotgun sequencing, we observe distinct microbiomes amongst two Atlantic cod (*Gadus morhua*) ecotypes (NEAC and NCC) with distinct behavior and habitats. In contrast, interspecific patterns of variation are more variable. For instance, we do not observe interspecific differentiation between the microbiomes of coastal cod (NCC) and Norway pout (*Trisopterus esmarkii*) whose lineages have evolutionary separated over 20 million years ago. The observed pattern of microbiome variation in these gadoid species is therefore most parsimoniously explained by differences in niche occupation.

## 1. Introduction

Significant research effort has focused on the importance of external, environmental factors (e.g. habitat, geography, microbial biodiversity, diet, water temperature or salinity) and internal, host-related factors (e.g. genetics, physiology or immunity) in driving the composition of the intestinal microbiome in fish (1, 2). That external factors play an important role is well established. For instance, bacterial diversity in the surrounding water influences the intestinal microbiome in fish larvae and fry (3, 4), water temperature is the main driver for the gut microbiome composition in farmed Tasmanian Atlantic salmon (*Salmo salar*) (5) and diet influences the intestinal composition in both experimental (6–9) as well as wild fish populations (10–13). Yet also internal factors influence the composition of these bacterial communities. For instance, observations of a shared (core) microbiome between wild and laboratory-raised zebrafish suggest that distinct selective pressures determine the composition of the microbial communities (14). Moreover, an association between host phylogeny and intestinal microbiome composition has been observed for a range of fishes, marine animals and terrestrial mammals (15–19).

The adaptive immune system appears especially important for host selection; individual variation of the Major Histocompatibility Complex (MHC) II correlates with the gut microbiome composition in stickleback (20); mucosal IgT depletion causes dysbiosis in rainbow trout (*Oncorhynchus mykiss*) (21), and lack of a functional adaptive immune system reduces the strength of host selection in knockout zebrafish models (22). Amongst bony fish, gadoid fishes have an unusual adaptive immune system –through the loss of MHC II, CD4 and invariant chain (Ii) and a range of innate (TLR) and MHC I immune-gene expansions (23, 24). Moreover, Atlantic cod has high levels of IgM (25) and a minimal antibody response after pathogen exposure (25–27). Gadoids therefore provide an interesting ecological system to study host-microbiome interactions (28).

Studies that specifically integrate internal and external influence support a role for both factors driving the microbial community composition (13, 29). Such studies however, remain restricted in both level of taxonomy of fishes (e.g. (30)) as well as taxonomical resolution of the microbial analyses (16S rRNA: 13, 29, 31–33). Importantly, it remains often difficult to separate the correlated effects of distinct behavior (i.e. diet) and niche occupation with interspecific selection and no comparative studies exist that use metagenomic shotgun sequencing to investigate fish populations with profound differences in behavior within a single species. It remains therefore unclear, whether the microbial composition for a range of wild fish species is characterized by intra- or interspecific divergence.

Here, we study intra- and interspecific divergence of intestinal microbial communities within the wide-spread family of *Gadidae* using a metagenomic shotgun dataset. We compare the microbiomes from Norway pout (*Trisopterus esmarkii*), poor cod (*Trisopterus minutus*), northern silvery pout (*Gadiculus thori*) and two ecotypes of Atlantic cod (*Gadus morhua*). These four species have overlapping geographical distributions, are dietary generalists, usually feeding over sandy and muddy bottoms on pelagic or benthic crustaceans, polychaetas and (small) fish (34, 35) and have evolutionary diverged approximately 20 million years ago (24). Norway pout is benthopelagic, distributed from the English Channel, around Iceland, and up to the Southwest Barents Sea, mostly found between 100 – 200 meters depth. Poor cod is also benthopelagic, distributed from the Trondheim Fjord in Norway to the Mediterranean Ocean, mostly found between 15 – 200 meters. Northern silvery pout (*Gadiculus thori*) is meso-to bathypelagic (36), distributed in the North Atlantic Ocean, along the coast of Norway and around Iceland and Greenland. It forms large schools, and are usually found between 200 – 400 meters (34, 36, 37). Finally, Atlantic cod has a trans-Atlantic distribution, from the Bay of Biscay to the Barents Sea, the Baltic Sea, around Iceland and Greenland, in the Hudson Bay and along the North American coast (34, 38–42). Atlantic cod comprises various subpopulations and “ecotypes” with distinct adaptations, migratory and feeding behavior. For instance, northeast Arctic cod (NEAC) performs typical spawning migrations from the Barents Sea to the Norwegian coast whereas the Norwegian coastal cod (NCC) remains more stationary (34, 43). These ecotypes have increased genomic divergence in several large chromosomal inversions (43–47), suggestive of local adaptation. The environments that these two ecotypes encounter are different, and they feed on distinct types of food. NEAC consumes mostly capelin and herring and NCC feeds on a wide range of crustaceans, fish and seaweed (34, 39, 48). During spawning, these ecotypes spatially co-occur, and long-term gene flow between ecotypes is supported by low overall estimates of divergence in most genomic regions, apart from the chromosomal rearrangements (43).

We hypothesize that if *interspecific selection* (indicative of host-selection) is the main driver for the intestinal communities in the *Gadidae*, most differences will be found *between the different species*, and not between the different ecotypes within Atlantic cod. In contrast, if *environmental factors* are the main drivers for the intestinal communities, we expect significant compositional differences *between the ecotypes* of Atlantic cod as well as varying levels of differentiation between the species. We use taxonomic profiling of metagenomic shotgun reads to classify these microbiomes –obtained from various locations around the Norwegian coast– at order and species-level resolution and analyze within-species differentiation of the most abundant members by genome-wide single nucleotide variation. Finally, differences in gut bacterial community composition among the species and ecotypes are assessed using multivariate statistics.

## 2. Methods

### 2.1 Sample collection

Northeast Atlantic cod (*Gadus morhua*) (NEAC, 10 individuals) were collected in Lofoten (N68.0619167, E13.5921667) in March 2014, and Norwegian coastal cod (*Gadus morhua*) (NCC, 10 individuals) at the same location in August 2014 (Fig. 1a, Table S1). NCC (2 individuals) were also collected in the Oslo Fjord (N58.9125100, E9.9202624 & N59.8150006, E10.5544914). Norway pout (*Trisopterus esmarkii*, 4 individuals), poor cod (*Trisopterus minutus*, 5 individuals) and northern silvery pout (*Gadiculus thori*, 3 individuals) were collected in the inner Oslo Fjord in May 2015 (Table S1). All fish specimens were collected from wild populations. A 3 cm long part of the hindgut (immediately above the short, wider rectal chamber) was aseptically removed *post-mortem* by scalpel and stored on 70% ethanol. The samples were frozen (−20°C) for long-term storage. Relevant metadata such as length, weight, sex and maturity were registered. We always strive to reduce the impact of our sampling needs on populations and individuals. Therefore, samples were obtained as a byproduct of conventional business practice. Specimens were caught by commercial vessels, euthanized by local fishermen and were intended for human consumption. Samples were taken post-mortem and no scientific experiments have been performed on live animals. This sampling follows the guidelines set by the “Norwegian consensus platform for replacement, reduction and refinement of animal experiments” (49) and does not fall under any specific legislation in Norway, requiring no formal ethics approval.

**Figure 1:**
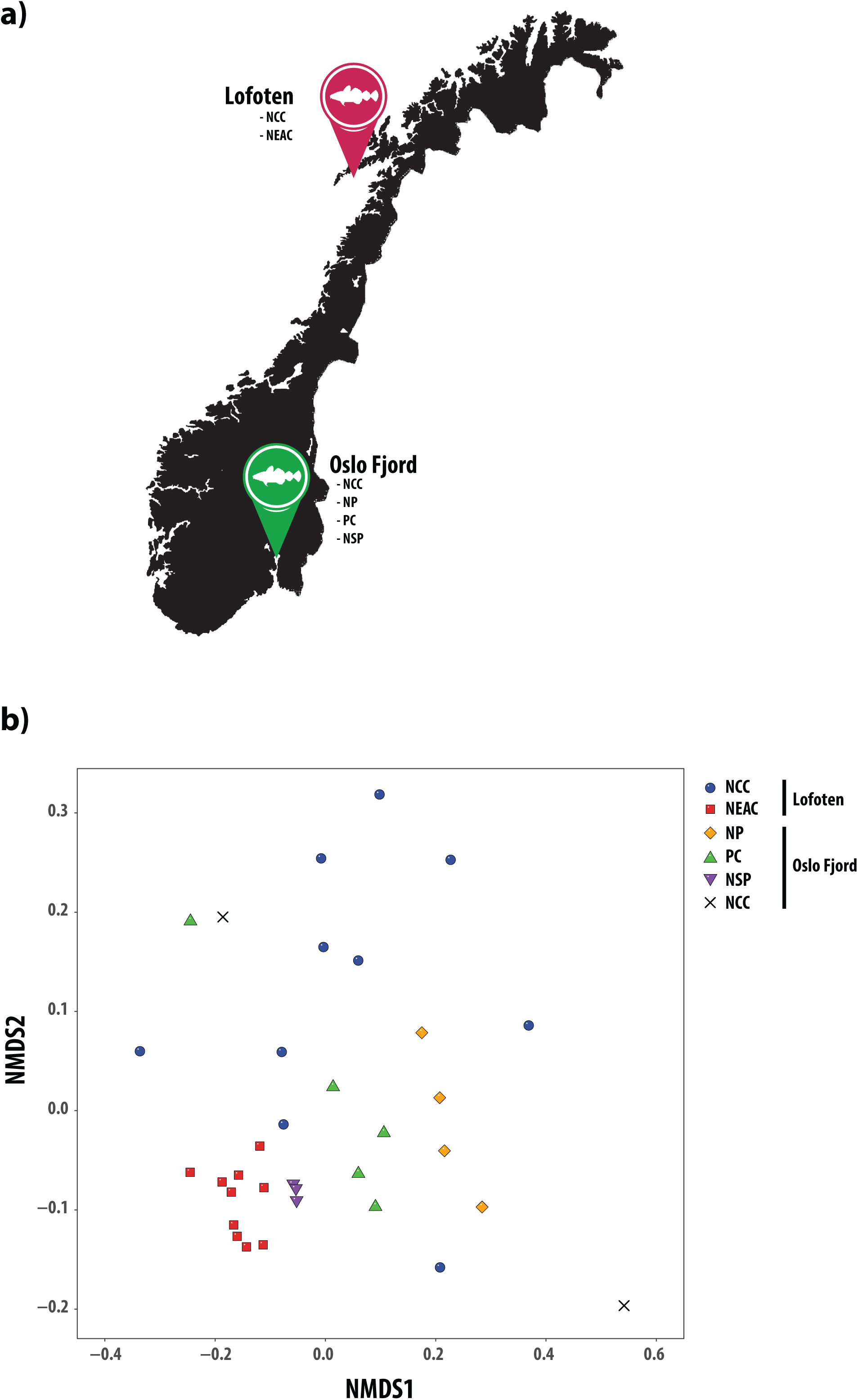
The intestinal microbiomes obtained from a range of gadoid species and ecotypes. **(A)** Map of sampling locations in Norway, Europe. Northeast Arctic cod (NEAC), and Norwegian coastal cod (NCC) were obtained from the Lofoten. NCC (two individuals), poor cod (PC), Norway pout (NP), and northern silvery pout (NSP) were obtained from the Oslo Fjord. **(B)** Non-metric multidimensional scaling (NMDS) plot of non-normalized, order-level sequence counts from the intestinal microbiomes of all samples. Each point represents an individual sample, and the species or ecotypes are indicated by different shapes and colors. The stress value of the NMDS plot is 0.14.

### 2.2 Sample preparation and DNA extraction

Intestinal samples were split open lengthwise, before the combined gut content and mucosa was gently removed using a sterile disposable spatula. Each individual sample was washed in 500 μl 100% EtOH and centrifuged before the ethanol was allowed to evaporate, after which dry weight was measured before proceeding to DNA extraction. DNA was extracted from between < 10 and 300 mg dry weight of gut content using the *MoBio Powersoil HTP 96 Soil DNA Isolation Kit* (Qiagen, Valencia, CA, USA) according to the DNA extraction protocol (v. 4.13) utilized by the Earth Microbiome Project (50). DNA was eluted in 100 μl Elution buffer, and stored at -20° Celsius. Due to high methodological consistency between biological replicates in previous experiments, only one sample was collected per fish (32).

### 2.3 Sequence data generation and filtering

Quality and quantity of the DNA was measured using a Qubit fluorometer (Life Technologies, Carlsbad, CA, USA), and normalized by dilution. DNA libraries were prepared using the *Kapa HyperPlus* kit (Roche Sequencing, Pleasanton, CA, USA) and paired-end sequenced (2×125 base pairs) on an Illumina HiSeq2500 using the HiSeq SBS V4 chemistry with dual-indexing in two independent sequencing runs. Read qualities were assessed using *FastQC* (51), before adapter removal, singleton read identification, de-duplication and further read quality trimming was performed using *Trimmomatic* (ver. 0.36) (52) and *PRINSEQ-lite* (ver. 0.20.4) (53) (Table S2). PhiX-, host- and human sequences were removed by mapping reads to the phiX reference genome [GenBank:J02482.1], the Atlantic cod genome assembly (gadMor 2) (this applied to all the fish species) (54), and a masked version of the human genome (HG19) (55) using *BWA* (ver. 0.7.13) (56) or *BBMap* (ver. 37.53) (JGI) with default parameters, and discarding matching sequences using *seqtk* (ver. 2012.11) (58). All sequence data have been deposited in the European Nucleotide Archive (ENA) under study accession number PRJEB31095.

### 2.4 Taxonomic profiling

Taxonomic classification of quality trimmed and filtered metagenomic paired-end reads was performed using *Kaiju* (ver. 1.5.0) (59) (“greedy” heuristic approach, -e 5), with the NCBI *nr* database (rel. 84) (incl. proteins from fungal and microbial eukaryotes) as reference (60). Counts of sequences successfully assigned to orders and species were imported into *RStudio* (ver. 1.1.383) (61) based on *R* (ver. 3.4.2) (62) for further processing. Filtering of the most abundant bacterial orders for visualization was based on a minimum relative abundance threshold of 1% of the total number of sequences per library (threshold ranging from 5933 - 95,146 depending on sample size). Similarly, filtering of the most abundant bacterial species was based on a minimum relative abundance threshold of 2% of the total number of sequences per library (threshold ranging from 6548 - 190,294 depending on sample size). Any taxon not exceeding this threshold in at least one (order level)/two (species level) sample(s) was removed. All filtering was based on the *R* package *genefilter* (ver. 1.62.0) (63). Final results were visualized using the R package *ggplot* (ver. 2.2.1) (64). Note: Based on a recent reclassification (65), we refer to the reference strain *Photobacterium phosphoreum* ANT-2200 (acc. nr. GCF_000613045.2) as *Photobacterium kishitanii* (Table S3).

### 2.5 Sequence variation analysis

In order to assess the heterogeneity of the most abundant bacteria in the fish species, we analyzed the sequence variation in the two genomes with the highest mean relative abundance over all fish species and ecotypes; *Photobacterium kishitanii* and *Photobacterium iliopiscarium*. Paired-end reads from each individual fish were mapped to the reference genomes (Table S3) using the *Snakemake* workflow (66) of *anvi’o* (ver. 5.1) (67) with default parameters in the “all-against-all” modus (with *anvi-profile --min-coverage-for-variability 20*). Samples of low coverage, restricting detection of SNVs in *anvi’o*, were excluded from the variation analysis. For each individual sample, variable sites were identified, and the mean number of these per 1000 bp calculated (variation density). A variable site required a minimum coverage of 20X. Next, variable sites with a minimum of 20X coverage in *all* samples were defined as single nucleotide variants (SNVs, *anvi-gen-variability-profile --min-occurrence 1 -- min-coverage-in-each-sample 20*). Coverage, variation density and SNV profiles were plotted in *RStudio* following the *R* script provided by *anvi’o* (68). The *anvi’o* SNV output was converted to .vcf format using a custom-developed script (https://github.com/srinidhi202/AnvioSNV_to_vcf), and the resulting .vcf files were used in a principal component analysis (PCA) to test for population differences as implemented in *smartpca* (ver. 6.1.4) (*EIGENSOFT*) (69).

### 2.6 Statistical analysis

Although included in data visualization, the Oslo Fjord NCC samples were excluded from statistical analysis, due to low sample size (n = 2). Within-sample diversity (alpha diversity) was calculated using the *diversity* function in the R package *vegan* (ver. 2.4-1) (70) based on Shannon, Simpson and Inverse Simpson indices calculated from non-normalized order-level read counts (Table S4). Differences in alpha diversity were studied using linear regression. The “optimal model” (the model that best describes the individual diversity) was identified through a “top-down” strategy including all covariates (Table S5), except weight, which highly correlated with length (r = 0.95), and selected through *t*-tests. Model assumptions were verified through plotting of residuals. Differences in bacterial community structure (beta diversity) between the fish species or ecotypes were visualized using non-metric multidimensional scaling (NMDS) plots based on the Bray-Curtis dissimilarity calculated from order-level sequence counts. Next, pairwise differences in beta diversity between the fish species or ecotypes were tested using Permutational Multivariate Analysis of Variance (PERMANOVA) in the *R* package *pairwise*.*adonis* (ver. 0.1) (71), a wrapper for the *adonis* functions in *vegan* (ver. 2.4-1), based on Bray-Curtis dissimilarity calculated from order-, genus- and species-level sequence counts. *pairwise*.*adonis* was run with 20,000 permutations, and *p* values adjusted for multiple testing using the Holm method (72). Adjusted *p*-values < 0.05 indicate statistical significance. PERMANOVA assumes the multivariate dispersion in the compared groups to be homogeneous; this was verified (*p* > 0.05) using the *betadisper* function (*vegan*) (Table S6). Similarity percentage (SIMPER) procedure implemented in *vegan* was used to quantify the contribution of individual orders to the overall Bray-Curtis dissimilarity between the species/ecotypes. All beta diversity analyses were based on sequence counts normalized using a common scaling procedure, following McMurdie & Holmes 2014 (73). This involves multiplying the sequence count of every unit (e.g. order) in a given library with a factor corresponding to the ratio of the smallest library size in the dataset to the library size of the sample in question. Normalization using this procedure effectively results in the library scaling by averaging an infinite number of repeated sub-samplings. We used Chi-squared statistics, as implemented in *smartpca* (69), to test for significant differences in the distribution of SNVs per reference genome, while correcting for multiple testing using sequential Bonferroni (72).

## 3. Results

### 3.1 Taxonomical composition of the intestinal microbiomes

We analyze a dataset of 422 million paired-end reads, with a median sample size of 11.9 million reads (8.0 - 19.6 million reads per sample) (Table 2, Table S7). Following filtering, order level classification could be obtained for 93% of all sequences (Table 2). Based on non-normalized order-level sequence counts, we observe clear patterns of separation between species and ecotypes in a multivariate NMDS plot (Fig. 1b), with NEAC and northern silvery pout forming distinct clusters, whereas the NCC populations encompasses the Norway pout and poor cod populations. *Vibrionales* is the most abundant order in the intestinal microbiomes of NCC specimens at both coastal locations (mean relative abundance (MRA): 76%) as well as Norway pout (MRA: 79%) and poor cod (MRA: 44%) (Fig. 2a, Table 3), with the remainder of each gut community consisting of a mix of orders with low relative abundance. The intestinal microbiome of the NEAC and northern silvery pout specimens have a significantly more diverse community composition (Fig. 2a, Fig. 3). NEAC is dominated by *Bacteroidales* (MRA: 21%), *Vibrionales* (MRA: 17%), *Clostridiales* (MRA: 12%) and *Brevinematales* (MRA: 7%) and northern silvery pout has a high relative abundance of orders *Brachyspirales* (MRA: 16%) and *Clostridiales* (MRA: 14%). Distinct from the gut community of the other fish populations, northern silvery pout has a low abundance of *Vibrionales*. Finally, the amount of sequences in the “Others” category, as well as sequences classified above order level (mean all samples: 7.8%), vary slightly between the fish species (Table S8). A species-level classification was obtained for 66% of all sequences. Overall, species of the genus *Photobacterium* comprise on average 40.6% of the classified sequences, ranging from 0.2% in northern silvery pout to 74.3% in Norway pout (Fig. 2b). In particular, *P. kishitanii* and *P. iliopiscarium* represent on average 43% and 36% of all *Photobacterium* species, although the ratio differs in the different fish species (e.g. 49% vs. 41% in NCC, 16% vs. 56% in NEAC and 55% vs. 12% in Norway pout).

**Table 1:** Species collected and sample location. The table lists the species, their Latin name, ecotype (for Atlantic cod), sampling location, number of specimens (*n*), and the abbreviation used in the current study.

| Species | Latin name | Ecotype | Sampling location | n | Abbreviation |
| --- | --- | --- | --- | --- | --- |
| Atlantic cod | *Gadus morhua* | Northeast Arctic cod | Lofoten | 10 | NEAC |
| Atlantic cod | *Gadus morhua* | Norwegian coastal cod | Lofoten | 10 | NCC |
| Atlantic cod | *Gadus morhua* | Norwegian coastal cod | Oslo fjord | 2 | NCC_Oslo |
| Poor cod | *Trisopterus minutus* |  | Oslo fjord | 5 | PC |
| Norway pout | *Trisopterus esmarkii* |  | Oslo fjord | 4 | NP |
| Northern silvery pout | *Gadiculus thori* |  | Oslo fjord | 3 | NSP |

**Table 2:** Overview of individual metagenomic sequence data from gadoid intestines. The table shows per sample the number of original reads, the percentage of reads remaining after trimming and filtering, percentage of host DNA, percentage of bacterial DNA and the final number of reads used in the microbiome analysis. PhiX- and human-derived DNA sequences represent a negligible proportion, and are excluded from the table. The bottom rows show total or mean values per column. On average, 42.2% of the quality filtered reads per sample are used for microbiome analysis. For details, see table S7.

| Sample | Raw reads | After quality<br>trimming/filtering (%) | Host DNA (%) | Bacterial DNA (%) | Final reads |
| --- | --- | --- | --- | --- | --- |
| NCC_01 | 10 883 740 | 85.9 | 87.3 | 12.7 | 1 187 649 |
| NCC_02 | 11 140 950 | 87.9 | 62.2 | 37.8 | 3 699 538 |
| NCC_03 | 9 891 322 | 90.2 | 41.2 | 58.8 | 5 249 515 |
| NCC_04 | 10 587 865 | 86.9 | 85.2 | 14.8 | 1 364 663 |
| NCC_05 | 8 423 091 | 89.1 | 57.7 | 42.3 | 3 171 737 |
| NCC_06 | 10 879 319 | 89.6 | 30.5 | 69.5 | 6 772 948 |
| NCC_07 | 10 082 237 | 91.8 | 31.3 | 68.7 | 6 361 506 |
| NCC_08 | 9 114 703 | 87.3 | 80.5 | 19.5 | 1 549 210 |
| NCC_09 | 11 105 189 | 89.1 | 62.2 | 37.8 | 3 733 846 |
| NCC_10 | 11 140 743 | 84.7 | 86.0 | 14.0 | 1 320 875 |
| NEAC_01 | 13 120 072 | 89.7 | 53.6 | 46.4 | 5 463 098 |
| NEAC_02 | 12 119 926 | 89.6 | 56.8 | 43.2 | 4 687 565 |
| NEAC_03 | 11 981 093 | 89.2 | 54.4 | 45.6 | 4 869 722 |
| NEAC_04 | 12 618 529 | 91.1 | 33.5 | 66.5 | 7 646 256 |
| NEAC_05 | 12 154 047 | 87.6 | 74.2 | 25.8 | 2 747 042 |
| NEAC_06 | 13 883 762 | 88.4 | 59.5 | 40.5 | 4 971 507 |
| NEAC_07 | 12 149 049 | 89.1 | 56.9 | 43.1 | 4 666 533 |
| NEAC_08 | 11 861 852 | 88.7 | 64.0 | 36.0 | 3 787 155 |
| NEAC_09 | 11 131 413 | 85.7 | 75.1 | 24.9 | 2 378 591 |
| NEAC_10 | 15 483 018 | 83.7 | 82.9 | 17.1 | 2 221 214 |
| OO_cod_01 | 8 047 125 | 83.7 | 85.9 | 14.1 | 949 039 |
| IO_cod_01 | 9 716 392 | 90.2 | 43.7 | 56.3 | 4 937 260 |
| NP_01 | 15 621 734 | 84.7 | 51.6 | 48.4 | 6 400 120 |
| NP_02 | 16 548 297 | 90.2 | 18.1 | 81.9 | 12 224 903 |
| NP_03 | 16 608 312 | 78.3 | 72.6 | 27.4 | 3 568 206 |
| NP_04 | 13 459 929 | 83.0 | 61.5 | 38.5 | 4 300 178 |
| PC_01 | 10 743 586 | 87.7 | 30.5 | 69.5 | 6 550 868 |
| PC_02 | 18 982 339 | 81.0 | 29.6 | 70.4 | 10 833 201 |
| PC_03 | 9 420 298 | 84.1 | 54.9 | 45.1 | 3 568 861 |
| PC_04 | 9 623 591 | 87.7 | 30.0 | 70.0 | 5 908 283 |
| PC_05 | 19 630 680 | 77.7 | 55.0 | 45.0 | 6 861 474 |
| NSP_01 | 14 283 994 | 73.2 | 67.5 | 32.5 | 3 396 962 |
| NSP_02 | 14 527 770 | 76.3 | 63.8 | 36.2 | 4 008 926 |
| NSP_03 | 15 261 446 | 80.9 | 66.6 | 33.4 | 4 123 131 |
| **Total:** | **422 227 413** |  |  |  | **155 481 582** |
| **Mean:** | **12 418 453** | **86.0** | **57.8** | **42.2** |  |

**Table 3:**
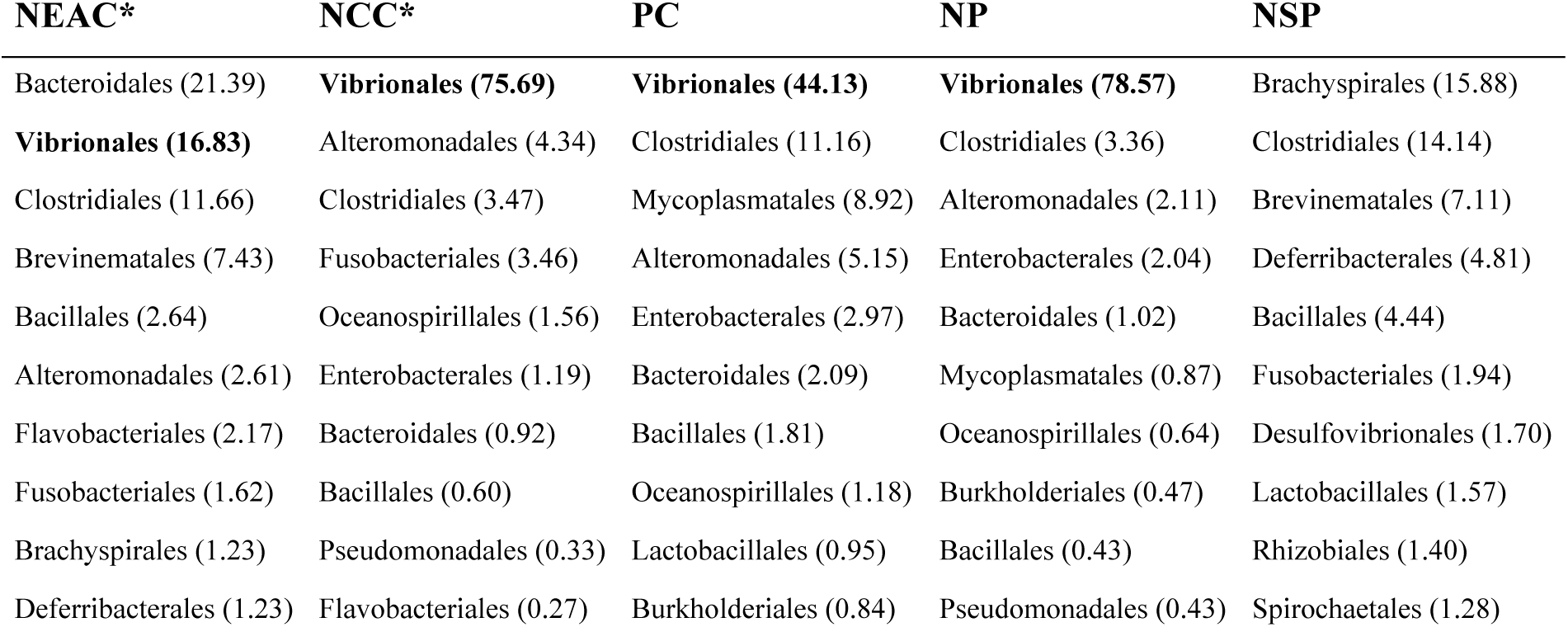
The 10 most abundant bacterial orders in the intestinal microbiomes of gadoid species and ecotypes. The table shows the mean relative abundance (%) of the ten most abundant orders in each of the species or ecotypes. *Vibrionales* is indicated in bold. The asterix indicates that these ecotypes belong to the same species (*Gadus morhua*)

**Figure 2:**
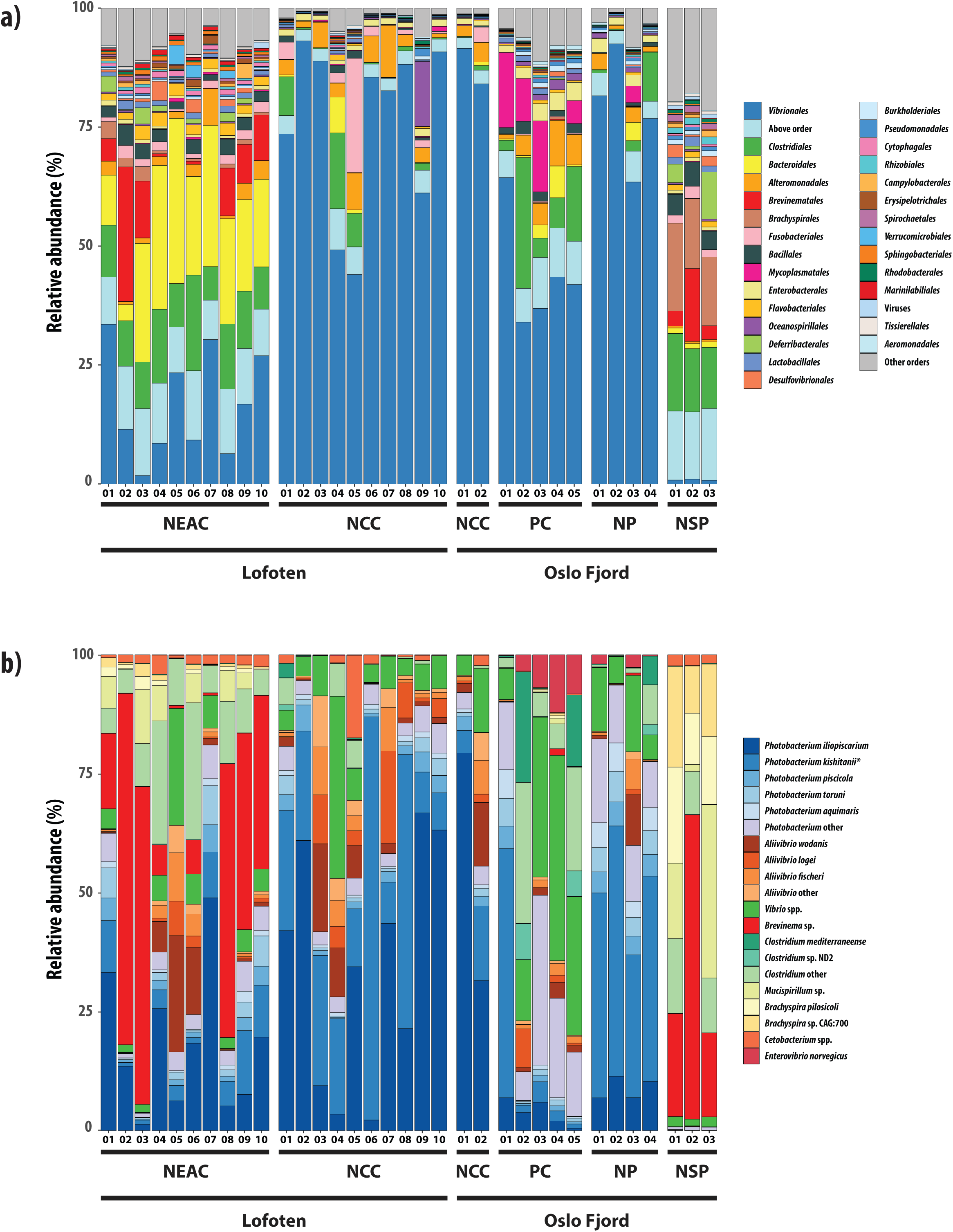
Taxonomic composition of the fish intestinal microbiomes. **(A)** Relative abundance of metagenomic shotgun sequences classified at the order level (93%). Colors represent the 28 orders with highest relative abundance, sequences assigned to other orders or viruses, and sequences classified above order level. Numbers along the x-axis indicate the individual samples of the different species/ecotypes. **(B)** Relative abundance of metagenomic shotgun sequences classified at the species level (66%). The plot includes the most highly abundant species, and other members of their parent bacterial genera (“other” categories) in the different fish species/ecotypes. Numbers along the x-axis indicate the individual samples of the different species/ecotypes. The star denotes the *P. kishitanii* species reclassified from *P. phosphoreum*.

**Figure 3:**
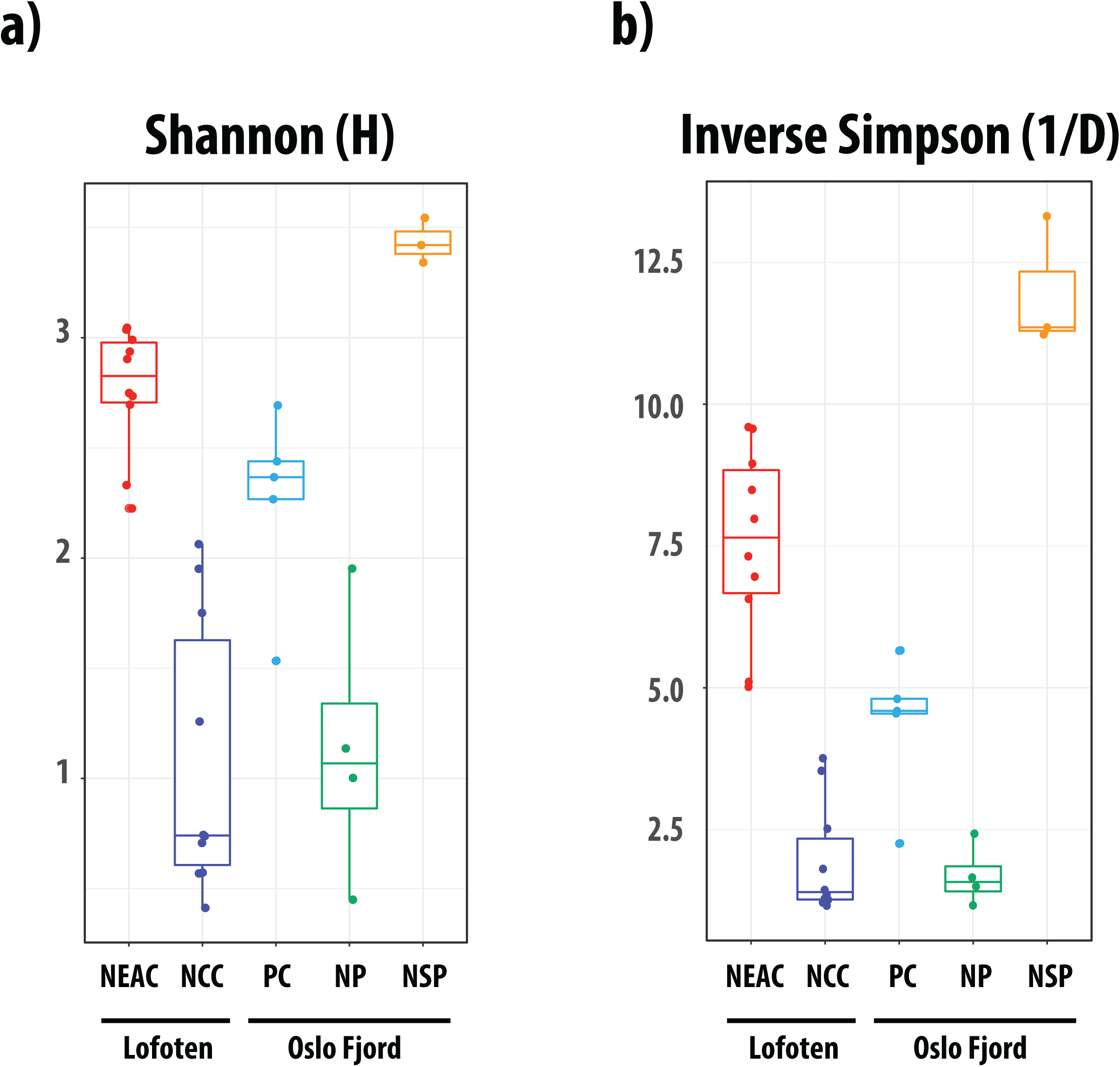
Within-sample microbial diversity in the gadoid species and ecotypes. Boxplots of Shannon **(A)** and Inverse Simpson **(B)** diversity in the fish species/ecotypes. Each individual is represented by a point, and the individuals are grouped and colored by species and ecotype. The middle band represents the median, while the upper and lower band shows the 75^th^ and 25^th^ percentile. The boxplots include the minimum and maximum alpha diversity values.

The NCC Lofoten intestinal microbiome is dominated by *P. iliopiscarium* (MRA: 21%) and *P. kishitanii* (MRA: 20%), followed by different species of *Aliivibrio* (*wodanis, logei* and *fischeri*) (MRA: 13%) (Fig. 2b). Similarly, the bacterial gut community of Norway pout is also dominated by *Photobacterium* species, in particular *P. kishitanii* (MRA: 17%). The intestinal microbiome of poor cod is dominated by *Photobacterium* species (MRA: 18%), followed by different *Vibrio* spp. (MRA: 8%). The gut bacterial community of NEAC is more diverse, with high relative abundance of a *Brevinema* sp. (MRA: 31%) and different species in the genera *Photobacterium* (MRA: 34%), *Clostridium* (MRA: 12%) and *Aliivibrio* (MRA: 9%). The high abundance of *Bacteroidales* observed at the order level (Fig. 2a) is not reflected at the species level, as this order represents a high number of *Bacteroidales* species with low abundance. Consequently, no *Bacteroidales* species are among the 15 most abundant species in the NEAC intestinal microbiome (Fig. 2b). The NEAC samples also contain a *Mucispirillum* sp. (MRA: 4%) and two *Brachyspira* spp. (MRA: 2%). In northern silvery pout, the gut microbiome is quite evenly distributed between the *Brevinema* sp., the *Mucispirillum* sp., *Brachyspira pilosicoli, Brachyspira* sp. CAG:700 and a group of different *Clostridium* species in two of three samples. The third sample contains the same species, but has an even higher relative abundance of the *Brevinema* sp. (64%) (Fig. 2b).

### 3.2 Variation in bacterial community composition among species and ecotypes

Significant differences in within-sample diversity (alpha diversity) at the order level are observed among all species and within-species ecotypes, except between NCC and Norway pout (Table 4, Table S5). None of the other covariates have a significant effect on alpha diversity. Similar to the results from the within-sample diversity, significant differences in community structure (beta diversity) are observed among the gadoid species at order-, genus- and species level (Table 5, Table S6). At the order level, the NEAC intestinal community has a different structure than what is observed in all the other gadoids (0.05 significance level). The NCC intestinal microbiome is also different from that of both poor cod and northern silvery pout. In agreement with results of within-sample (alpha) diversity, no differences in community structure are observed between the microbiomes of NCC and Norway pout. Finally, no differences are observed between the gut microbiome of poor cod vs. Norway pout, poor cod vs. northern silvery pout or Norway pout vs. northern silvery pout (*p* = 0.074 for all). Beta diversity analysis also demonstrate that community differences at the genus and species level are similar to those observed at the order level (Table S6).

**Table 4:** Effects of covariates on the intestinal microbial diversity (alpha diversity) of gadoid species and ecotypes. Results from the optimal linear regression models used in testing for significant effects of covariates on within-sample (alpha) diversity based on non-normalized, order-level sequence counts. Population (species/ecotype) is the only covariate with a significant effect, and estimates are given relative to NCC. Significant effects (*p* < 0.05) are indicated in bold.

|  | Shannon |  | Simpson |  | Inv. Simpson |  |
| --- | --- | --- | --- | --- | --- | --- |
|  | Estimate | p-value | Estimate | p-value | Estimate | p-value |
| Intercept | 1.08 | **0.0000** | 0.38 | **0.0000** | 1.93 | **0.0001** |
| NEAC | 1.69 | **0.0000** | 0.48 | **0.0000** | 5.62 | **0.0000** |
| NP | 0.06 | 0.8367 | -0.01 | 0.8784 | -0.25 | 0.7476 |
| PC | 1.18 | **0.0001** | 0.37 | **0.0002** | 2.44 | **0.0017** |
| NSP | 2.36 | **0.0000** | 0.53 | **0.0000** | 10.03 | **0.0000** |

**Table 5:**
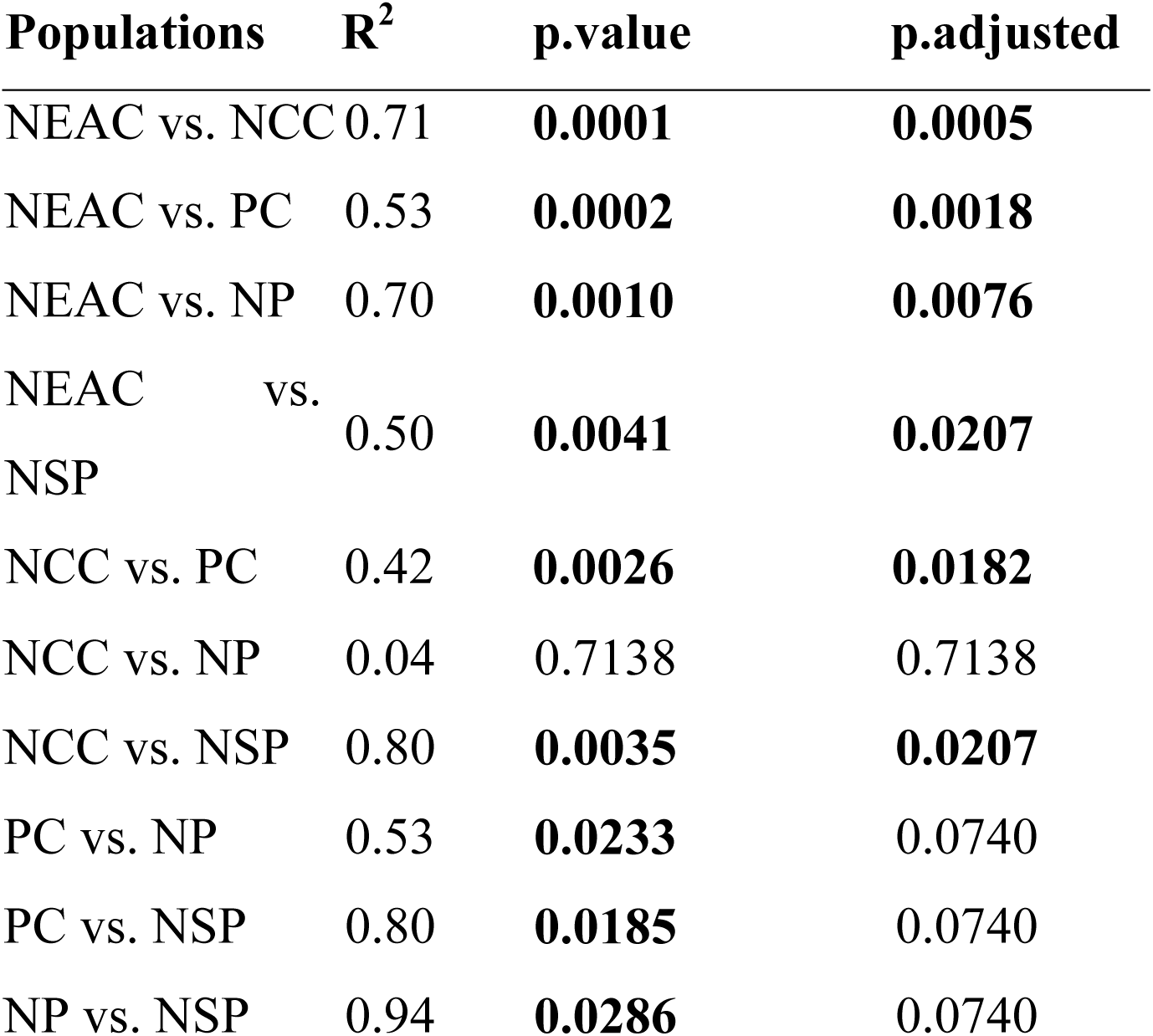
PERMANOVA analysis of intestinal microbial diversity from gadoid species and ecotypes. The table shows R^2^ values, *p*-values and adjusted *p*-values for pairwise comparisons of community composition (beta diversity) between the different species or ecotypes using PERMANOVA. The tests are based on Bray-Curtis dissimilarity calculated from order-level, normalized sequence counts. *p*-values are adjusted for multiple testing by the Holm method. Significant differences (*p* < 0.05) are indicated in bold. Genus- and species-level results are shown in Table S6.

| Populations | $R^2$ | p.value | p.adjusted |
| --- | --- | --- | --- |
| NEAC vs. NCC | 0.71 | **0.0001** | **0.0005** |
| NEAC vs. PC | 0.53 | **0.0002** | **0.0018** |
| NEAC vs. NP | 0.70 | **0.0010** | **0.0076** |
| NEAC vs. NSP | 0.50 | **0.0041** | **0.0207** |
| NCC vs. PC | 0.42 | **0.0026** | **0.0182** |
| NCC vs. NP | 0.04 | 0.7138 | 0.7138 |
| NCC vs. NSP | 0.80 | **0.0035** | **0.0207** |
| PC vs. NP | 0.53 | **0.0233** | 0.0740 |
| PC vs. NSP | 0.80 | **0.0185** | 0.0740 |
| NP vs. NSP | 0.94 | **0.0286** | 0.0740 |

Differences in the intestinal community composition between these gadoids are predominantly explained by changes in the relative abundance of a limited number of orders. For example, different proportions of *Vibrionales* contribute 29% to the (Bray-Curtis) dissimilarity between the NCC and NEAC (*p* = 0.001), followed by differences in the relative abundance of *Bacteroidales*, explaining 10% of the dissimilarity (*p* = 0.001) (Table S9). Together, 80% of the observed dissimilarity between NCC and NEAC is explained by differences in their relative abundance of the top six orders. Similarly, 60% of the dissimilarity between NCC and northern silvery pout are driven by *Vibrionales, Brachyspirales* and *Clostridiales*.

### 3.3 Bacterial within-species variation of Single Nucleotide Variant heterogeneity

We investigated bacterial within-species variation of *P. iliopiscarium* and *P. kishitanii* –with sufficient read coverage across all samples– among the different gadoids by mapping sequencing reads to their respective reference genomes (GCF_000949935.1, GCF_000613045.2). In the samples used for SNV analysis, the mean percentage of the reference genomes with minimum 20-fold coverage (coverage breadth) after mapping were 63% for *P. iliopiscarium* and 19% for *P. kishitanii*. Hence, the variation analysis of the two species is based on different proportions of the reference genomes. The two reference genomes greatly vary in the number of SNVs observed in all samples, from 84,866 in *P. iliopiscarium* to 1229 in *P. kishitanii*, Fig. 4a). The density of variable sites within each individual sample shows varying levels of heterogeneity in the bacterial populations (Fig. 4b). This heterogeneity is particularly clear in *P. kishitanii*, with sites-density varying from 0.5 to 45.4 variant positions per Kbp per individual specimen. Further, the heatmap shows gadoid specific SNV patterns (Fig. 4c), in particular for *P. iliopiscarium*, where Norway pout contains a distinct pattern compared to the other gadoids, indicating the presence of specific *P. iliopiscarium* strain(s). Statistical analyses of SNV variation reveals that NEAC has a significantly different SNV pattern from Norway pout (Chi-square, *p* = 0.017) and poor cod (*p* = 0.028) for *P. kishitanii*, and from NCC (*p* = 0.033) and Norway pout (*p* = 0.000) for *P. iliopiscarium* (Fig. 4d, Table S10). NCC has a significantly different SNV pattern from Norway pout (*p* = 0.003) for *P. iliopiscarium*. (Fig. 4d, Table S10). The relative abundance of *P. kishitanii* and *P. iliopiscarium* vary greatly among the fish specimens used in the variation analysis (Fig. 4e).

**Figure 4:**
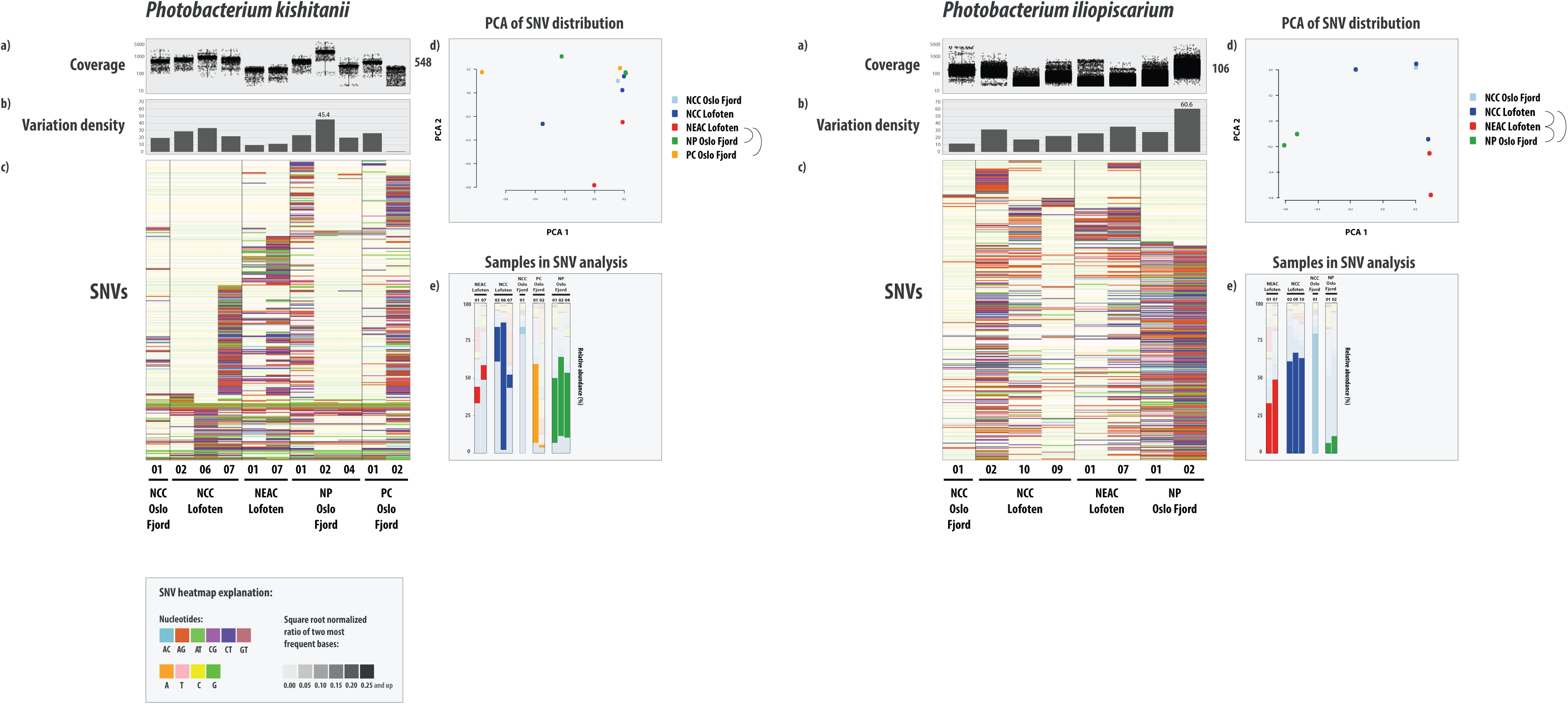
SNV variation analysis of the most abundant bacterial genomes in the microbiomes of gadoid species. For both of the genomes, the figure displays **(A)** read coverage per single nucleotide variant (SNV) position in each sample from the different species/ecotypes (mean coverage on right side of plot), **(B)** variation density (number of variable positions per 1000 kbp. reported in each individual sample, independent of coverage in the other samples) per sample (maximum value indicated). The y-axis of the coverage- and variation density plots are scaled across the genomes. **(C)** Heatmap of a randomly chosen subset of 400 SNVs. In the heatmap, each row represents a unique variable nucleotide position, where the color of each tile represents the two most frequent competing nucleotides in that position. The shade of each tile represents the square root-normalized ratio of the most frequent two bases at that position (i.e., the more variation in a nucleotide position, the less pale the tile is); see legend in the bottom of the figure. **(D)** Principal component analysis (PCA) plot of the SNV distribution (within-species variation) among the different samples. Each sample is represented by a dot, and colored according to species or ecotype membership. Half-circles to the right of the legend indicate species or ecotypes with significantly different within-species variation (i.e. different strains). **(E)** Relative abundance of the different samples used in variation analysis. The bars are colored according to the SNV plot in (D).

## 4. Discussion

Using metagenomic shotgun sequencing, we show the composition of the intestinal microbiomes of two Atlantic cod ecotypes (NEAC, NCC) to be at least as divergent as those found between the different codfish species investigated here. Our findings have several implications for our understanding of the composition of the intestinal microbiome in wild fish populations.

Although species-specific selection has been proposed as a factor driving the composition of the intestinal community in fish in a variety of settings (13, 14, 16–19, 29, 33), our results show that this may not be the most important driver among gadoid species in wild populations. First, we observe highly significant differences in the intestinal microbiomes at *order*-, *species*- and *within-species* bacterial level between the NEAC and NCC ecotypes. Despite showing different migratory behavior, these ecotypes co-occur during seasonal spawning in northern Norway (Lofoten), from where most of the samples are collected (43–45). Second, we observe no significant bacterial *order-* or *species-*level differences in the intestinal microbiome between different gadoid species, Atlantic cod (ecotype NCC) and Norway pout, which are sampled from different geographical locations (Lofoten and Oslo fjord). We visually do not observe any differentiation between the NCC sampled from Lofoten and the Oslo fjord (although statistical certainly is low), which reflects an earlier observed lack of geographical structure for this ecotype (30). The similar microbial composition of the NCC and Norway pout is striking, as these are distinctly different genetic lineages with an evolutionary separation of at least 20 million years (24). These results suggest that NCC and Norway pout occupy an environmental niche that allows bacterial members with a broad geographical distribution to colonize their intestinal communities. Overall, the observation of a significant differentiation between microbiomes from ecotypes of the same species *and* a lack of differentiation between microbiomes from two distinct species, suggest that the intestinal microbiome in these gadoid species and ecotypes is not driven by species-specific selection alone.

There are several factors that may underlie the compositional differences in the NCC and NEAC intestinal microbiomes. First, for more than 10 months during the year, the two populations encounter different habitats, as the NEAC ecotype is distributed in the pelagic waters of the Barents Sea, while NCC remains more stationary in coastal waters (74). Although several 16S rRNA-based studies have reported limited effects of geographic location on the composition and diversity of the fish intestinal microbiome (32, 75), the Barents Sea has significantly lower temperatures (76) compared to Norwegian coastal waters (77). Temperature has been shown to have a significant impact on the intestinal microbiome in several studies (e.g., Senegalese sole (*Solea senegalesis*), Tasmanian Atlantic Salmon (*Salmo salar*) and mummichog (*Fundulus heteroclitus*) (5, 78, 79) but not in all cases (e.g., Atlantic salmon (80). Second, the ecotypes were sampled during different seasons; NCC Lofoten during summer (August) and NEAC during winter/early spring (March). Nonetheless, a lack of difference between NCC Lofoten (August) and NCC Oslo fjord (May) suggests that seasonality is unlikely to fully explain the observed differences between NEAC and NCC. Third, the ecotypes show different feeding behavior; while the NEAC during foraging and spawning migrations from the Barents Sea may perform vertical movements down to 500 meters (42, 81, 82), NCC mainly occupy shallow and warmer coastal and fjord waters (83). This behavior exposes the two ecotypes to different sources of food, with NEAC predominantly eating capelin and herring (48), and NCC living of a more diverse diet, including crustaceans, fish and even seaweeds (34, 39). Diet has been shown to influence the composition of the intestinal microbiome in several fish species (9, 10, 13, 78, 84, 85). Finally, Barents Sea has a high microbial biodiversity compared to coastal areas (86). The specific bacterial load in the surrounding waters also influences the intestinal microbiome composition in fish, including Atlantic cod (3, 4). Nonetheless, because these different environmental and behavioral factors are correlated, it is unclear which of these parameters contributes the most to the observed differences in the intestinal microbiome composition between these ecotypes.

Comparing two spatially separated coastal Atlantic cod populations, metagenomic shotgun data revealed no strain-level differentiation (30). In this study, we find specific SNV variants amongst the most abundant bacterial species that are associated with either species or Atlantic cod ecotype. This indicates that NEAC harbor different strains of *P. iliopiscarium* than those identified in the NCC ecotype and the other gadoid species. Our current study encompasses a significantly greater geographical area and taxonomical samples than the earlier coastal comparison (30–32), and is indicative of strain-level variation at such larger comparative scales. In line with Riiser et al. 2019 (30), this study shows that such strain-level differences cannot be detected using 16S rRNA techniques alone, and that metagenomic shotgun sequencing is currently the most accurate approach to detect strain-level spatial variation in the marine environment.

Most striking amongst the comparisons of gadoid species are the microbiome differences observed in NEAC, northern silvery pout and poor cod compared to NCC and Norway pout. Several bacterial species that drive this differentiation are of particular interest. First, two bacterial species, *Mucispirillum* sp. and *Brevinema* sp., are almost exclusively detected in the intestinal microbiomes of NEAC and northern silvery pout. Nonetheless, these genera are represented by a single species in the *RefSeq* database (60) (accessed 10.01.19) and hence little is known. *B. andersonii* (order *Brevinematales*) was originally identified in short-tailed shrews (*Blarina brevicauda*) and white-footed mice (*Peromyscus leucopus*), and were found unable to grow below 25°C (87). *Brevinema* sp. has previously been identified in Atlantic cod (32) and in Atlantic salmon (88). *Mucispirillum schaedleri* (order *Deferribacterales*) is a mucosa-associated member of the intestinal microbiome in terrestrial animals as pigs, goats and rodents, where it is thought to be involved in mucus production through expression of lectins, important components in the innate immune response (89, 90). Nevertheless, the distant relationship between Atlantic cod and these terrestrial hosts, and the availability of only single reference genomes for *Mucispirillum* and *Brevinema*, strongly suggests that the representatives found here represent related, but novel species with a different intestinal ecology and physiology. Second, both NEAC and northern silvery pout contain significant fractions of *Brachyspira* spp., previously identified as dominant members in the gut of the carnivorous marine fish species mahi mahi (*Coryphaena hippurus*) (12, 91). *Brachyspira* spp. are known as intestinal pathogens in pigs and humans (92, 93), although recent studies show that *Brachyspira* spp. are more widespread in the wildlife community than previously thought, including in freshwater (94). The ecology of *Brachyspira* in the marine environment is unclear, although an association with the carnivorous diet of mahi mahi and NEAC may suggest that the diet of northern silvery pout also has a considerable carnivorous component. Third, poor cod is the only species with considerable abundance of *Enterovibrio norvegicus* (Table S11). This bacterium within the *Vibrionaceae* family was isolated from the intestines of cultured turbot (*Scophthalmus maximus*) larvae in Norway, and classified as a novel species phenotypically similar to the *Vibrio* genus (95). Interestingly, poor cod are also host to the highest abundance of *Vibrio* spp. among the fish species in this study (Table S11). Other *Enterovibrio* species have been found in association with diseased corals (96) and internal organs of cultured fish species in the Mediterranean Ocean (97–99). However, little is known about the function of this relatively novel genus in fish intestines.

Given the observations of species specific selection for a similar microbiome in various teleosts and range of habitats (13, 14, 16–19, 29, 33), the diverse microbiomes *within* and *among* gadoid species may suggest that their intestinal communities could be more easily modulated by external factors. At this stage, limited sampling across various fish taxa and the lack of comparative approaches leave reasons for such diverse communities speculative. Nonetheless, it is interesting to note that all gadoids have unusual adaptive immune system –through the loss of MHC II, CD4 and invariant chain (Ii) and a range of innate (TLR) and MHC I immune-gene expansions (23, 24). There are significant correlations between immune genes and the vertebrate microbiome (100, 101), and it has been hypothesized that adaptive immunity has evolved to help maintain complex community of beneficial commensal bacteria (102). Indeed, studies of wild-type zebrafish and knockout zebrafish without a functional adaptive immune system suggested that adaptive immunity increases the strength of host filtering of potential fish-associated microbes (22). The unusual adaptive immune system of gadoids may therefore affect the strength of co-evolutionary associations with their microbiome.

## 5. Conclusion

Based on metagenomic shotgun sequencing, we here characterize the intra- and interspecific community composition among two ecotypes of Atlantic cod and three related fish species in the *Gadidae* family. Several of these fish species harbor unique, and possibly novel bacterial species. We identify a complex pattern of diversity with significant differences between the Atlantic cod ecotypes, and variable interspecific patterns of variation. Although most species and ecotypes yield different communities, those found in coastal cod (NCC) and Norway pout are not significantly diverged, indicating that ecological niche plays an important role in determining the intestinal microbiomes in these gadoid species.

## Authors’ contributions

SJ, BS and THA conceived and designed the experiments. KSJ provided the initial framework for the study. ESR and SJ sampled the specimens. ESR performed the laboratory work. ESR and THA performed data analysis. SV created the Python script to convert the anvi’o format to VCF. ØB, THA, ESR and BS interpreted the results. ESR and BS wrote the paper with input of all authors. All authors read and approved the final manuscript.

## Acknowledgements

We thank Børge Iversen and Helle Tessand Baalsrud for their kind help in sampling Atlantic cod specimens in Lofoten, and Martin Malmstrøm, Paul Ragnar Berg and Monica Hongrø Solbakken for sampling at Sørøya. We are grateful for the metagenome sequencing performed at the Norwegian Sequencing Centre (NSC: https://www.sequencing.uio.no).

## Availability of data and materials

The data set generated and analyzed for this study is available in the European Nucleotide Archive (ENA), study accession number PRJEB31095.

